# Effective Tyrosine Kinase Inhibitors result in the intracellular accumulation of EGFR and allows response prediction in patients

**DOI:** 10.1101/798314

**Authors:** Maurice de Wit, Ya Gao, Darlene Mercieca, Iris de Heer, Bart Valkenburg, Martin van Royen, Joachim Aerts, Peter Sillevis Smitt, Pim French

## Abstract

Clinical responses to EGFR tyrosine kinase inhibitors are restricted only to tumors harboring specific activating mutations and even then, not all tyrosine kinase inhibitors provide clinical benefit. We here show that the addition of EGFR-TKIs results in a strong and rapid intracellular accumulation of the protein. However, this accumulation was observed only in the context of a combination of a TKI-sensitive mutation with a clinically effective TKI: TKI-insensitive mutations did not show this accumulation nor did clinically ineffective TKIs induce accumulation. All TKIs effectively inhibited EGFR phosphorylation and downstream pathway activation, irrespective of the mutation present in EGFR. The discrepancy between molecular activity of TKIs and their efficacy in patients therefore is mimicked by the mutation- and TKI-specificity of intracellular accumulation. Using this intracellular accumulation as assay, we were able to predict response to gefitinib in a panel of cell-lines (harboring different EGFR mutations) and predicted clinical benefit to EGFR TKIs on a cohort of unselected pulmonary adenocarcinoma patients (hazard ratio 0.21, P=0.0004). Even in patients harboring rare mutations with unknown TKI-sensitivity, intracellular accumulation was predictive of the clinical response. The intracellular accumulation depended on a continued presence of TKI indicating that TKIs exert a continued effect on the protein even after its dephosphorylation. It is therefore possible that accumulation is caused by conformational changes induced by both the mutation and the TKI and this change induces a block in intracellular trafficking. Interestingly, intracellular accumulation was observed independent of the genetic background of the cell, indicating that accumulation is almost entirely dictated by the combination of mutation and TKI. Our results therefore suggest that TKI-sensitivity is tumor-type independent.

## Introduction

The epidermal growth factor receptor (EGFR) gene is a key oncogene that is mutated in many different cancer types including gliomas, colorectal cancer and pulmonary adenocarcinoma. Tumors depend on EGFR signaling for their growth and this dependency makes EGFR an attractive target for therapy. Indeed, pulmonary adenocarcinoma patients harboring EGFR mutations show strong clinical response to EGFR tyrosine kinase inhibitors (TKIs) [1-4]. Unfortunately, other tumor types that depend on EGFR signaling, such as glioblastomas (the most common and aggressive type of primary brain cancer), show no response to EGFR-TKIs [5-7].

Not all EGFR-mutated pulmonary adenocarcinoma patients benefit from EGFR TKIs: responses are predominantly observed in the context of deletions in exon 19 or missense mutations L858R, G719X and S768I. Patients with other, less common-activating mutations such as exon 20 insertions show no benefit from EGFR TKIs (see e.g. mycancergenome.org) despite EGFR being effectively dephosphorylated [8-10]. Apart from this mutation-specificity, there also a drug-specificity of clinical responses: where several EGFR-TKIs (erlotinib, gefitinib, afatinib, dacomitinib and osimertinib) have provided clinical benefit to *EGFR*-mutated pulmonary adenocarcinoma patients, a phase II study on lapatinib did not show any sign of clinical activity [10-12]. This lack of clinical activity is surprising as lapatinib is a highly potent inhibitor of EGFR phosphorylation. In summary, clinical responses to EGFR TKIs is restricted to a limited set of mutations only, and not all TKIs are clinically effective. The molecular mechanisms for this mutation- and drug-specificity remains unknown.

We here describe a simple in-vitro assay that can predict which mutation is sensitive to which TKI. Similar to the responses observed in the clinic, our assay is both mutation and TKI-specific, and is independent on the inhibition of EGFR-phosphorylation and downstream pathway activation. We validate our assay on eleven different cell-lines that harbor as many different EGFR mutations and in a variety of unselected EGFR-mutated pulmonary adenocarcinoma patients. We also predict responses in patients harboring EGFR mutations where TKI sensitivity thus-far has not been documented. The observed TKI-induced intracellular accumulation is likely a result of a block in intracellular trafficking due to a continued association of the TKI with EGFR. Because the intracellular accumulation was observed independent of the genetic background of the cell, accumulation is almost entirely dictated by the combination of mutation and TKI and this independence argues that all patients with sensitive EGFR mutations should, regardless of the type of tumor, be considered for treatment with EGFR-TKIs.

## Results

### Clinically effective TKIs induce an intracellular accumulation of EGFR

To examine mutation- and TKI-specificity of clinical responses, we generated eGFP-tagged EGFR mutation constructs, stably expressed them in HeLa cells and monitored response to inhibitors *in-vitro*. When erlotinib was addition to cells expressing EGFR^L858R^, we observed a striking intracellular accumulation of the protein visible as intracellular EGFR-protein ‘spots’ (dozens per cell and up to thousands per imaging field, figure 1a). Using an automated quantitative imaging analysis setup, we show that the response was dose dependent, occurred almost immediately following drug administration and persisted for >3 days (figure 1b/c and supplementary figure 1 and supplementary movie 1). In contrast, erlotinib did not induce the intracellular accumulation in cells expressing EGFR-wildtype or EGFRvIII (a deletion of exons 2-7, the most common mutation in GBMs, figure 1a) but did in a construct containing both EGFRvIII and L858R mutation (EGFR^v/8^, not shown) demonstrating that the accumulation is mutation dependent. Moreover, the intracellular accumulation was observed in cells expressing EGFR^L858R^ only after the addition of clinically effective drugs erlotinib, gefitinib, dacomitinib or osimertinib but not after administration of lapatinib, a drug that does not show clinical efficacy (figure 1b/c/d). The intracellular accumulation of EGFR in our assay therefore was mutation and TKI-dependent.

**Figure 1:**
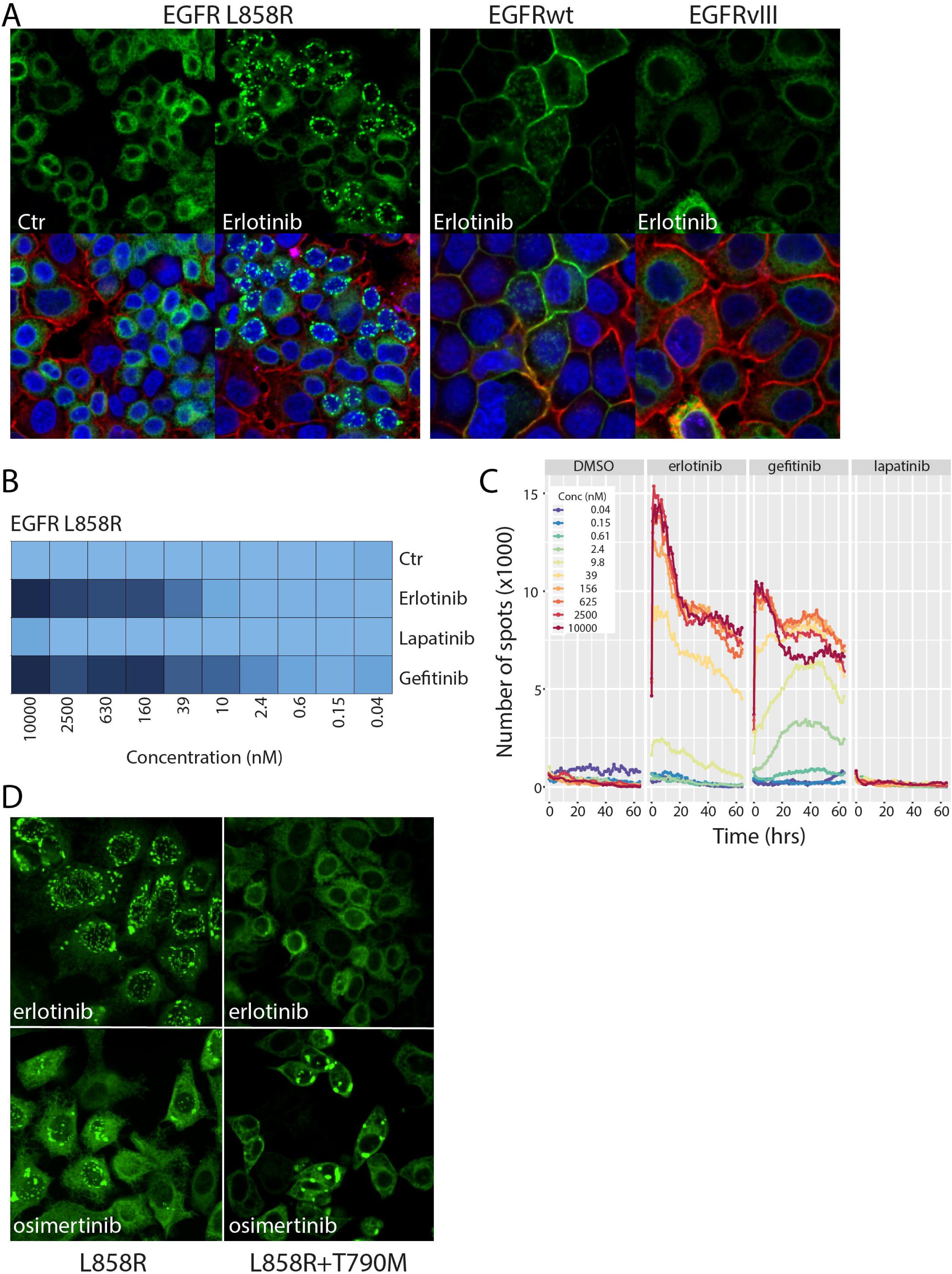
Clinically effective EGFR TKIs induce a rapid and massive intracellular accumulation of EGFR. **A)** Erlotinib treatment of HeLa cells ectopically expressing EGFR^L858R^ results in its intracellular accumulation. This accumulation is not observed in cells expressing EGFRwt or EGFRvIII. Top panels depict the EGFR signal only (Green); bottom panels is a merge including Red: WGA (membrane) and blue: Hoechst (nucleus). **B)** intracellular accumulation is dose dependent and only occurs with clinically effective inhibitors erlotinib and gefitinib but not with lapatinib. The intracellular accumulation is retained up to 60 hrs **(C). D)** Erlotinib no longer induces intracellular accumulation in cells ectopically expressing the resistance mutation EGFR^L858R+T790M^. They do however remain responsive to osimertinib (bottom panels).

Osimertinib is a potent third generation EGFR inhibitor with clinical activity also in tumours harboring the secondary T790M resistance mutation [3]. A construct harboring this T790M (EGFR^L858R+T790M^) secondary resistance mutation no longer responded to erlotinib or gefitinib in our assay, but strongly induced intracellular accumulation following addition osimertinib (figure 1d). Constructs harboring secondary resistance mutations therefore only induced intracellular accumulation in response to a TKI that is clinically effective on this mutation.

HeLa cells were chosen as model for these initial experiments as they do not depend on EGFR for their growth and neither inhibitors nor the intracellular accumulation induced death in these cells (not shown). This simple model system therefore avoids potential confounding effects of cell death and associated mechanisms and focusses on the (direct) effects inhibitors have on EGFR. Accumulation was however not specific to HeLa cells as erlotinib, gefitinib and osimertinib but not lapatinib, strongly induced the intracellular accumulation in U87 cells expressing EGFR^L858R^ but not in cells expressing EGFRvIII or EGFRwt (supplementary figure 2). We also created stable cell lines in which non-tagged EGFR was expressed from a bicistronic EGFR-IRES-eGFP vector. Similar to the eGFP-tagged mutation constructs, effective EGFR TKIs erlotinib, gefitinib, dacomitinib and osimertinib, but not lapatinib, led to the intracellular accumulation of EGFR, but only in EGFR^L858R^-IRES-eGFP expressing cells and not in EGFRwt-IRES-eGFP expressing cells (figure 2). These data confirm our observation that clinically effective EGFR-TKIs result in the accumulation of intracellular EGFR but only in the context of TKI-sensitive mutations.

**Figure 2:**
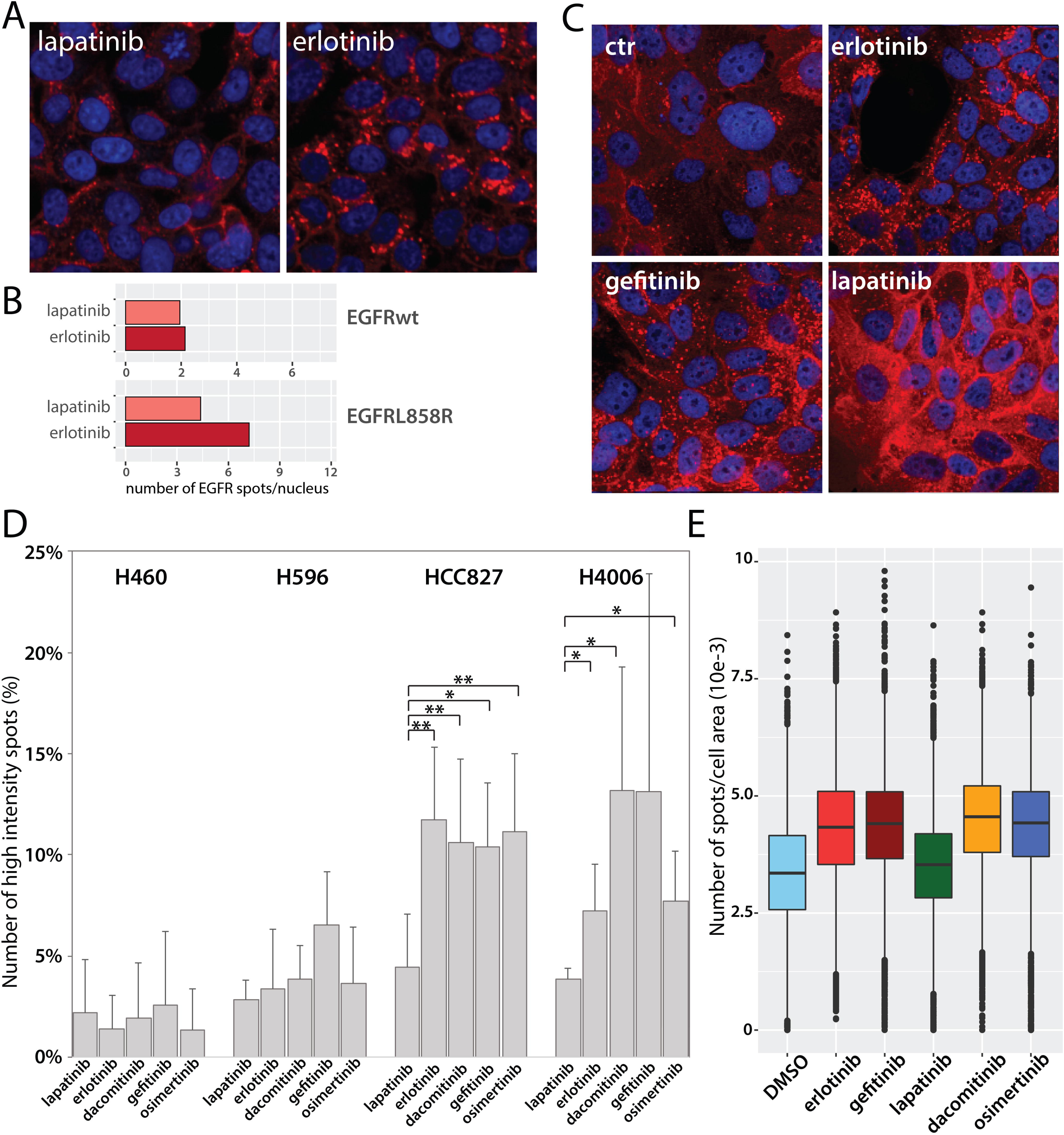
Intracellular accumulation of untagged EGFR and in lung cancer cell lines. A) erlotinib but not lapatinib induces intracellular accumulation of EGFR in HeLa cells expressing EGFR^L858R^-IRES-eGFP. B) quantification of images in A showing lapatinib induces accumulation only in EGFR^L858R^-IRES-eGFP expressing cells (lower graph) and not in EGFRwt-IRES-eGFP expressing cells (top graph). C) Also the HCC827 lung cancer cell line (containing a TKI sensitive mutation), erlotinib and gefinitnib, but not lapatinib induced intracellular accumulation of EGFR. Quantification of images shown in C demonstrates that both the number of high-intensity spots (D) and the total number of spots (E) increase following treatment with erlotinib, gefitinib, dacomitinib or osimertinib, and not by lapatinib, but only in cell lines harbouring TKI-sensitive mutations (HCC827 and H4006).

To further evaluate intracellular EGFR accumulation, we used four different lung cancer cell lines that harbour endogenous EGFR mutations. Although all four lung cancer cell lines tested had relatively high numbers of EGFR-positive intracellular vesicles at baseline, also in these cell lines a significant increase in the intracellular accumulation of EGFR was observed when cells were incubated with clinically effective TKIs (erlotinib, gefinitnib, dacomitinib and osimertinib) but not by the clinically ineffective TKI lapatinib (figure 2). This increase in lung cancer cell lines was observed as an increase in the number of EGFR-positive intracellular vesicles and in the intensity thereof (n = 5 independent replicates). The inhibitor-induced intracellular accumulation was only observed in cell lines harbouring TKI-sensitive mutations (HCC827 and H4006) and not in cell lines that do not harbour TKI-sensitive mutations (H596 nor H460). Effective EGFR TKIs therefore lead to the intracellular accumulation of EGFR, also in cells harbouring endogenous EGFR mutations.

### Intracellular accumulation predicts response to gefitinib in cell lines

Because of the correlation of the intracellular accumulation with responses observed in the clinic, we tested whether intracellular accumulation was able to actually predict response to EGFR TKIs. For this, we screened the Genomics of Drug Sensitivity in Cancer (GDSC) database that contains drug-sensitivity data in >1000 genomically characterized cell-lines [13-15]). We selected 11 cell lines with a known EGFR mutation (10 different mutations) with documented response to gefitinib. We then generated constructs for all EGFR mutations, stably expressed them in HeLa cells and screened for inhibitor-induced intracellular accumulation. Constructs EGFR^L858R^, EGFR^E746_A750del^, EGFR^L747_E749del^, EGFR^S768I^, and EGFR^G719S^ all responded to gefitinib by rapidly inducing intracellular accumulation of EGFR; none of the other mutation constructs showed such accumulation (supplementary figure 3). Dose response analysis indicated that EGFR^L858R^ and EGFR^E746_A750del^ were highly sensitive to gefitinib (IC50 <20 nM) whereas EGFR^L747_E749del^, EGFR^S768I^ and EGFR^G719S^ showed considerably higher IC50 values (156, 625 and 456 nM respectively, figure 3).

**Figure 3:**
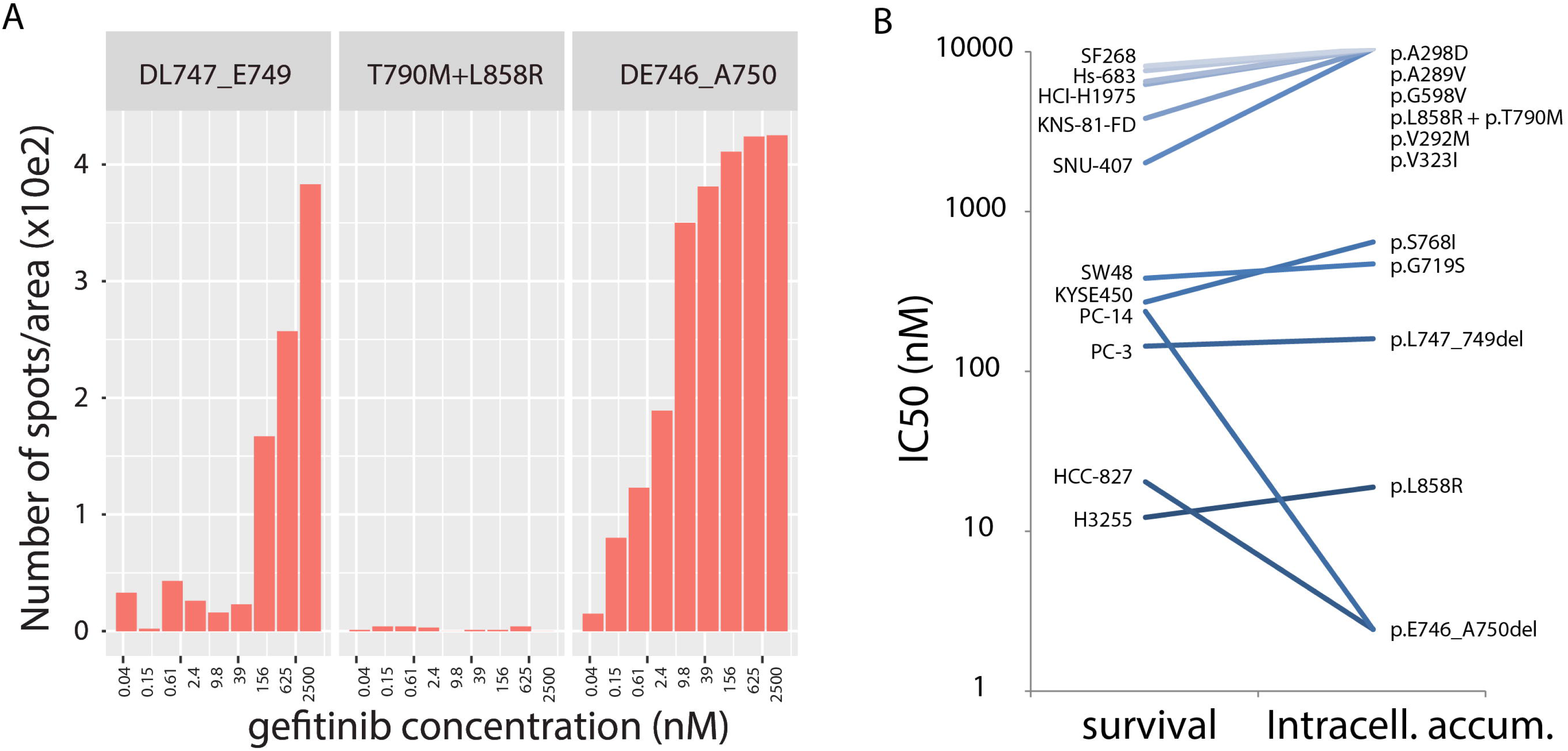
Intracellular accumulation of EGFR formation predicts sensitivity to gefitinib in GDSC cell lines. **A)** Examples of dose response analysis of intracellular EGFR accumulation. As can be seen, cells expressing EGFR^E746_A750del^ have a high sensitivity to accumulate EGFR compared to cells expressing EGFR^L747-E749del^. No intracellular accumulation is observed in cells expressing EGFR^L858R+T790M^. B) Comparison between the ability of gefitinib to induce intracellular accumulation in HeLa cells (IC50 value for intracellular accumulation) with the sensitivity to gefitinib in the EGFR-mutated GDSC cell-lines (i.e. the IC50 value for viability, see also supplementary table 1). Despite differences in the cell-lines and assays used, we find a high concordance between cell viability and inhibitor-induced EGFR intracellular accumulation.

Comparing ‘*gefitinib induced intracellular accumulation in HeLa cells expressing EGFR-mutation constructs*’ with ‘*gefitinib sensitivity of cells endogenously expressing EGFR mutations*’ showed that the IC50 value for intracellular accumulation was highly similar to the IC50 value for viability (extracted from the GDSC database, supplementary table 1) for each of the mutations tested. Cell lines that are highly sensitive to gefitinib also harbored mutations that were highly sensitive to gefitinib induced intracellular accumulation (EGFR^L858R^ or EGFR^E746_A750del^), cell-lines with moderate sensitivity harbored mutations that were moderately sensitive to gefitinib induced intracellular accumulation (EGFR^L747_E749del^, EGFR^S768I^ or EGFR^G719S^) and cell-lines that are insensitive to gefitinib harbored mutations that do not show gefitinib induced intracellular accumulation (figure 3). Of note, virtually identical results were obtained using erlotinib in our assay and lapatinib was unable to induce intracellular accumulation in any mutation construct. Our relatively simple and straightforward assay therefore was able to predict sensitivity to EGFR TKIs in cell lines harboring endogenous EGFR mutations.

**Table 1.**
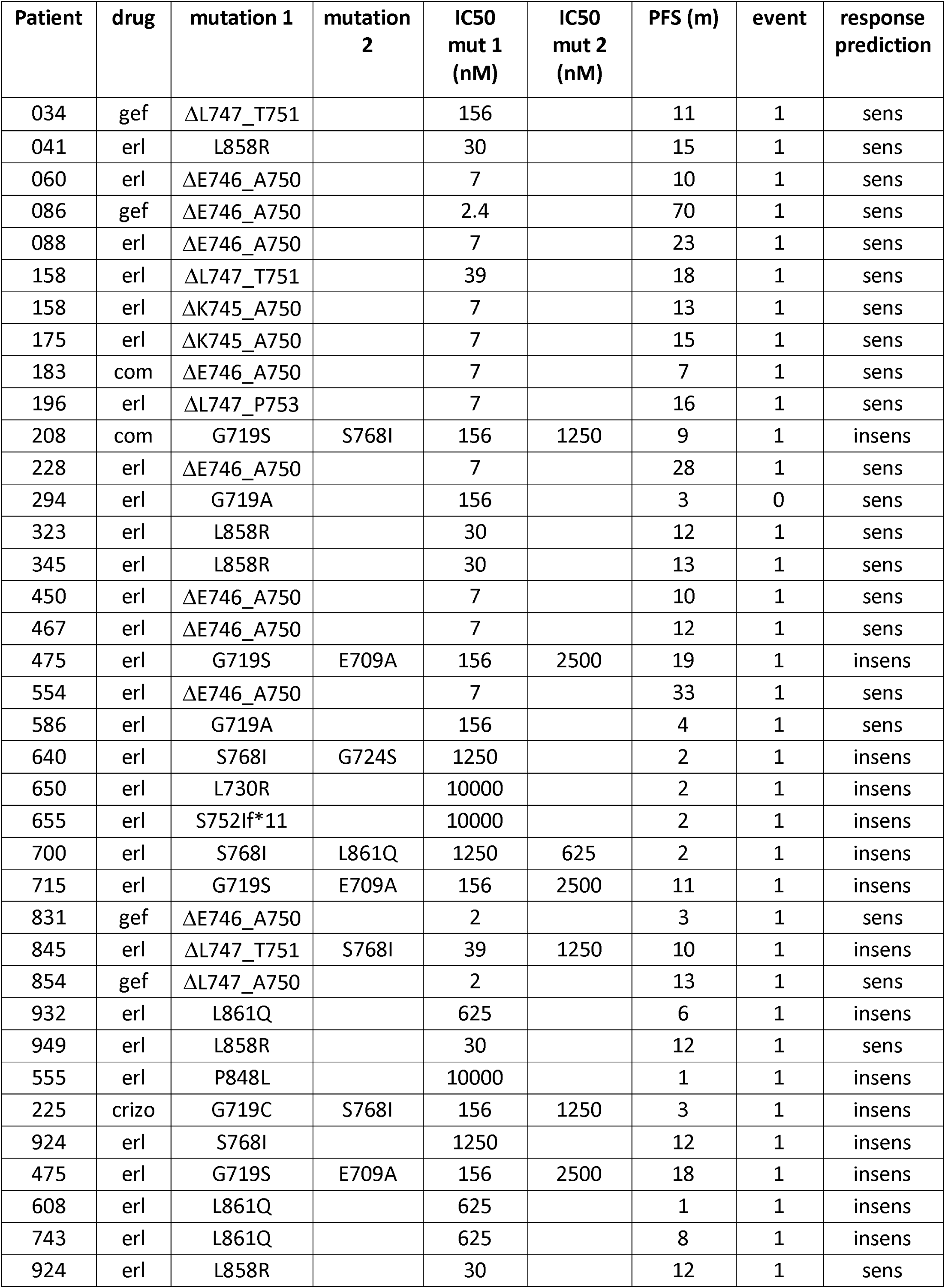
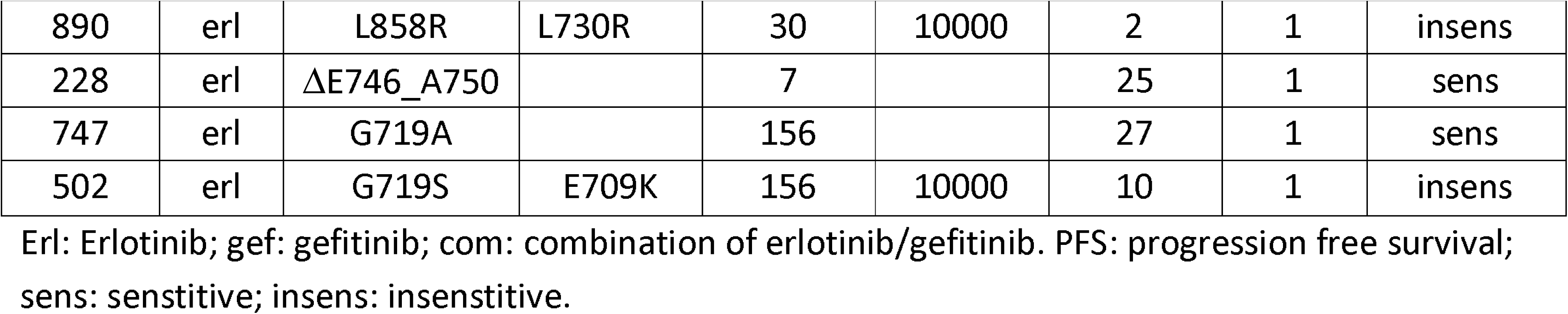
Intracellular accumulation formation predicts response to EGFR TKIs in pulmonary adenocarcinoma patients

### Intracellular accumulation predicts response to EGFR TKIs in pulmonary adenocarcinoma patients

To determine whether intracellular accumulation of EGFR can predict response to TKIs in patients, we screened all pulmonary adenocarcinoma patients treated in 2016 and 2017 within our clinic for the presence of EGFR mutations (table 1). For each mutation identified in this unselected patient cohort, we generated EGFR-mutation constructs and stably expressed them in HeLa cells. In each EGFR mutation we tested the ability of TKIs to induce intracellular accumulation and, if so, determined the IC50 value thereof. All experiments were performed using automated image analysis software and were blinded to clinical outcome. We then split the dataset into ‘predicted responders’ and ‘predicted non-responders’ using a cutoff of 500 nM for intracellular EGFR accumulation. This cutoff was defined prior to performing the experiments and was based on estimates of the intra-tumoral concentration of erlotinib (∼200ng/g tumor tissue, though there is a wide inter-patient and intra-tumoral variability [16]). On this dataset, we show that ‘predicted responders’ had a significantly longer time to progression to first line EGFR TKIs than the ‘predicted non-responders’ (median survival 7.0 vs 13 months, HR 0.21, P=0.0004, figure 4). These data demonstrate that intracellular accumulation of EGFR s predictive for clinical response to first line EGFR TKI.

**Figure 4:**
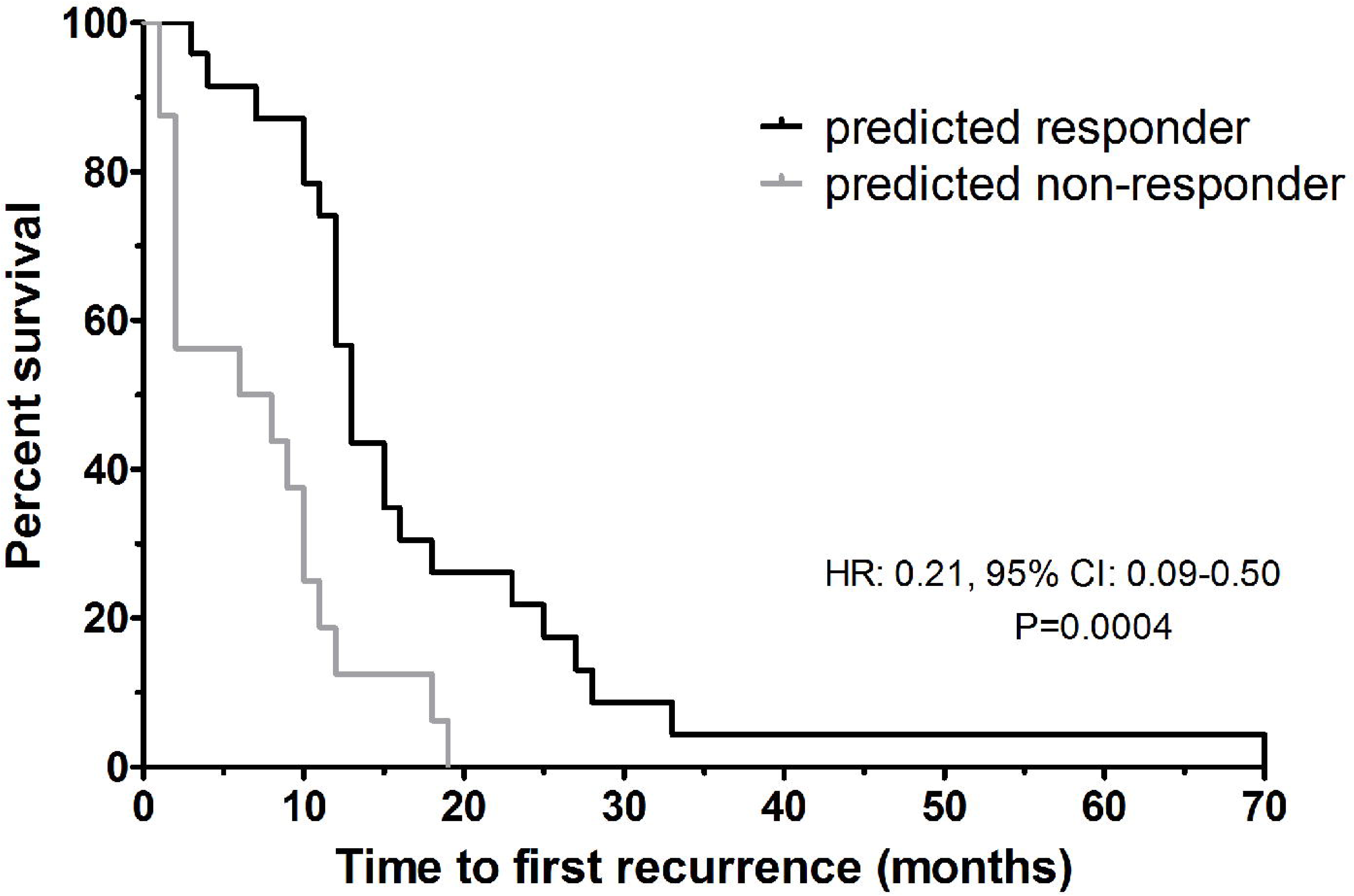
Intracellular accumulation of EGFR predicts response to first line treatment in pulmonary adenocarcinoma patients. Mutation constructs for all activating mutations in table 1 (n=41) were generated and the IC50 value for intracellular accumulation to various TKIs was determined. Patients were then separated into predicted responders and non-responders (blinded to clinical outcome using a predefined cutoff of 500 nM, i.e. a clinically achievable concentration). As can be seen, intracellular accumulation predicts progression free survival in response to first line TKI treatment in pulmonary adenocarcinoma patients.

### Predicting response to rare, unknown mutations

We further evaluated the intracellular accumulation in mutations where clinical responses to EGFR TKIs is unknown. Because of the rarity of such mutations, we included DIRECT database queries and public domain literature to assess clinical responses (table 2). The EGFR^T751-I759delinsATA^ mutation showed strong intracellular accumulation (IC50 for gefitinib and erlotinib of 40 and 10 nM respectively) and was classified as ‘predicted responder’. A patient with similar mutation indeed showed a partial response to EGFR TKIs and a progression free survival of 8 months [17]. The EGFR^L747-E749del^ showed sufficient strong intracellular accumulation (IC50 for gefitinib and erlotinib of 156 and 432 nM respectively) to be classified as ‘predicted responder’. The DIRECT database identified two patients harboring such mutations and both showed partial responses to EGFR TKIs (PFS 6 months in one patient, PFS not reported for the other) [18]. The EGFR^E746A^ missense mutation did not show any sign intracellular accumulation and was classified as ‘predicted non-responder’. Two patients have been described harboring a similar mutation and neither patient responded to EGFR TKI treatment (both had stable disease, no PFS reported) [19, 20]. Finally, the EGFR^P848L^ was found in one of our patients and, as predicted by a lack of intracellular accumulation, this patient did not respond to EGFR TKI treatment. A patient with identical mutation also did not respond to erlotinib [21]. Therefore, also in these novel mutations with previously unknown sensitivity to EGFR-TKIs, intracellular EGFR accumulation highly correlated to the clinical responses in all seven patients. These results therefore further demonstrate that intracellular accumulation predicts response to EGFR TKIs.

**Table 2.** Response prediction of unknown EGFR mutations

| **Mutation** | **Response prediction** | **Clinical response** | **PFS** | **ref** |
| --- | --- | --- | --- | --- |
| p.L747_E749del | sensitive | PR | 6 | Yeh et al., 2013 |
| p.L747_E749del | sensitive | PR |  | Yeh et al., 2013 |
| p.E746X | insensitive | SD |  | Kalikaki et al., 2010 |
| p.E746X | insensitive | SD |  | Pallis et al., 2007 |
| P848L | insensitive |  | 1 | this manuscript |
| P848L | insensitive | SD | 4.6 | Faehling et al., 2017 |
| p.T751_I759del | sensitive | PR | 8 | Schrock et al., 2016 |

PR: partial response; SD: stable disease.

### All EGFR TKIs effectively inhibit EGFR and its pathway

Because of the strong phenotype induced by effective EGFR TKIs, but only on TKI-sensitive mutations, we explored whether these TKIs and/or mutations differ with respect to pathway activation and inhibition. Western blot analysis showed that all inhibitors effectively blocked EGFR phosphorylation in HCC827 cells (that contains an endogenous EGFR^E746-A750del^ mutation, figure 5a). In a cell line containing the T790M resistance mutation (H1975), only osimertinib reduced EGFR phosphorylation (supplementary figure 4a). Two other lung cancer cell-lines (H460 and H596, EGFR wt and amplified respectively), showed no EGFR phosphorylation under normal serum culture conditions (supplementary figure 4a, see also [22, 23]). Quantitative image analysis, using pan- and phospho-specific EGFR antibody stainings, confirmed the efficacy of EGFR-TKIs: In cell lines containing activating EGFR mutations (HCC827 and HCC4006), EGFR is phosphorylated and the addition of all tested TKI effectively inhibited this phosphorylation (figure 5b/c). In cell lines without activating EGFR mutations (NCI-H460 and H596), EGFR is not phosphorylated and EGF stimulation resulted in a rapid increase in EGFR phosphorylation levels. Addition of EGFR TKIs prior to EGF stimulation prevented EGFR-phosphorylation and the addition of TKIs after EGF stimulation resulted in a rapid dephosphorylation of EGFR (figure 5d/e, supplementary figure 4b). Also in stably transfected HeLa cells, all intracellular accumulation consisted of dephosphorylated EGFR (supplementary figure 5). All TKIs therefore effectively block EGFR phosphorylation and therefore cannot explain the differences in the observed intracellular accumulation.

**Figure 5:**
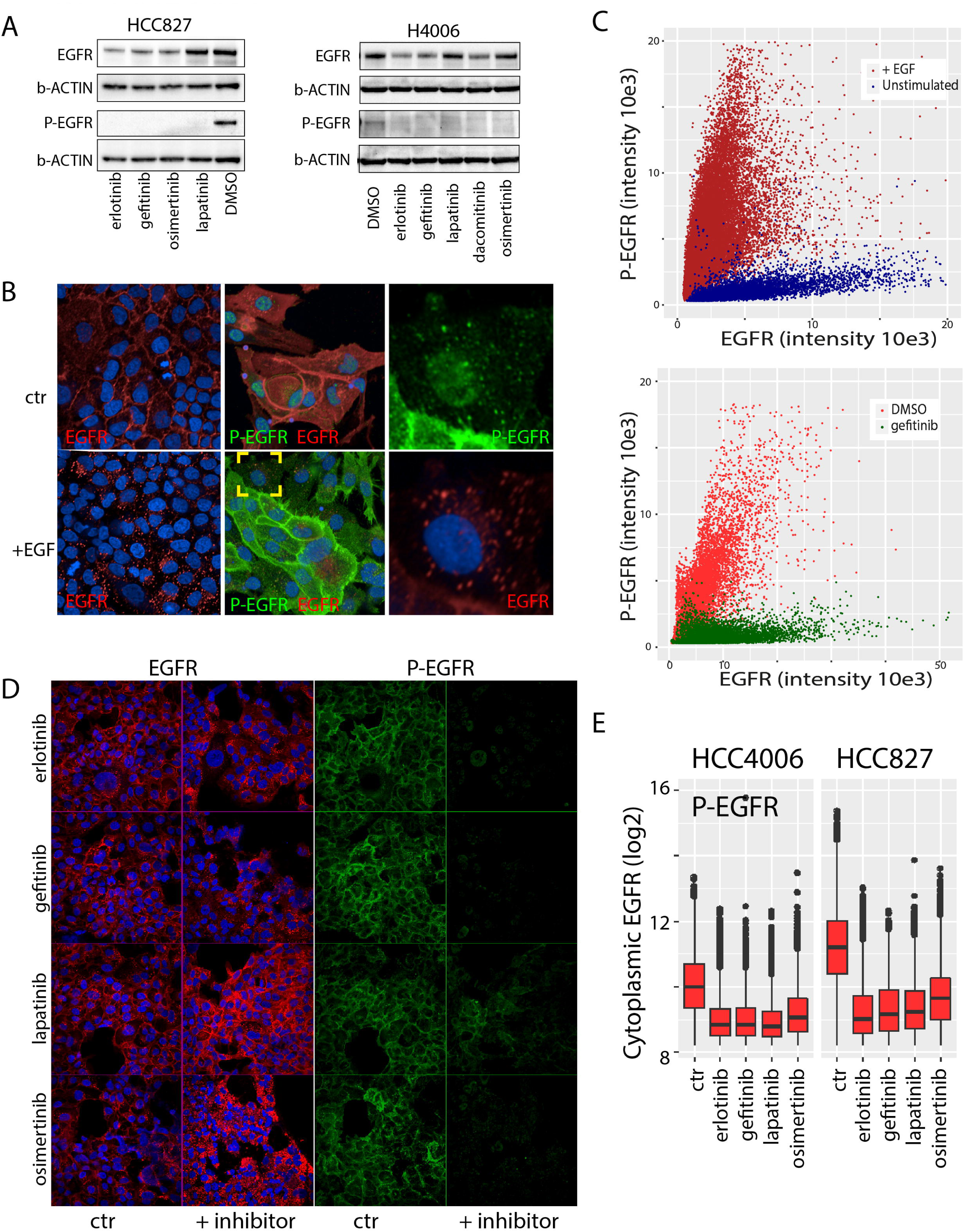
All TKIs effectively inhibit EGFR phosphorylation. **A)** all EGFR TKIs effectively block EGFR phosphorylation on western blot in HCC827 and H4006 cell-lines. **B)** imaging analysis showing effects of EGF stimulation on EGFR (and EGFR phosphorylation) in H460 (left panels) and H596 cells (middle and right panels). In H460 cells, EGF stimulation results in internalization of the receptor. Co-staining for phospho-EGFR shows a rapid increase in EGFR-phosphorylation, which overlaps with the pan-EGFR signal. Right panels are an inset of the yellow square in EGF-stimulated H596 cells, depicting phospho-EGFR staining (top) and pan-EGFR staining (red). **C**) Quantification of the phospho-EGFR signal in areas staining for pan-EGFR. As can be seen, EGF stimulation of H596 cells (top panel) results in a very pronounced increase in phospho-EGFR staining per cell (each dot represents an individual area that stained positive for EGFR). In cells HCC827 cells (lower panel) that have constitutive active EGFR phosphorylation, gefitinib significantly decreases the phospho-EGFR signal. **D)** HCC827 cells stained for EGFR (red, left panels) and phospho-EGFR (green, right panels). As can be seen, all inhibitors effectively reduce EGFR phosphorylation. **E)** quantification of images presented in D) as presented in C).

We performed reversed phase phosphoprotein arrays (RPPA) to study whether different TKIs and/or mutations differentially affect pathway activation. We find that erlotinib and lapatinib are equally effective in blocking downstream EGFR signaling (figure 6a-c, supplementary table 2) irrespective of the type of EGFR mutation present and irrespective of the inhibitor used: in all three cell lines tested phosphorylation of AKT (serine 473), mTOR (serine 2448) and P90 (threonine 573) was inhibited by the addition of erlotinib or lapatinib. We also did not identify differences in other molecular pathways interrogated by the RPPA arrays between the two inhibitors. RT-qPCR further demonstrated that EGFR-TKIs effectively blocked the expression the immediate early genes *EGR1* and *cFOS*, also irrespective of EGFR mutation type or inhibitor used [24-26] (figure 6d).

**Figure 6:**
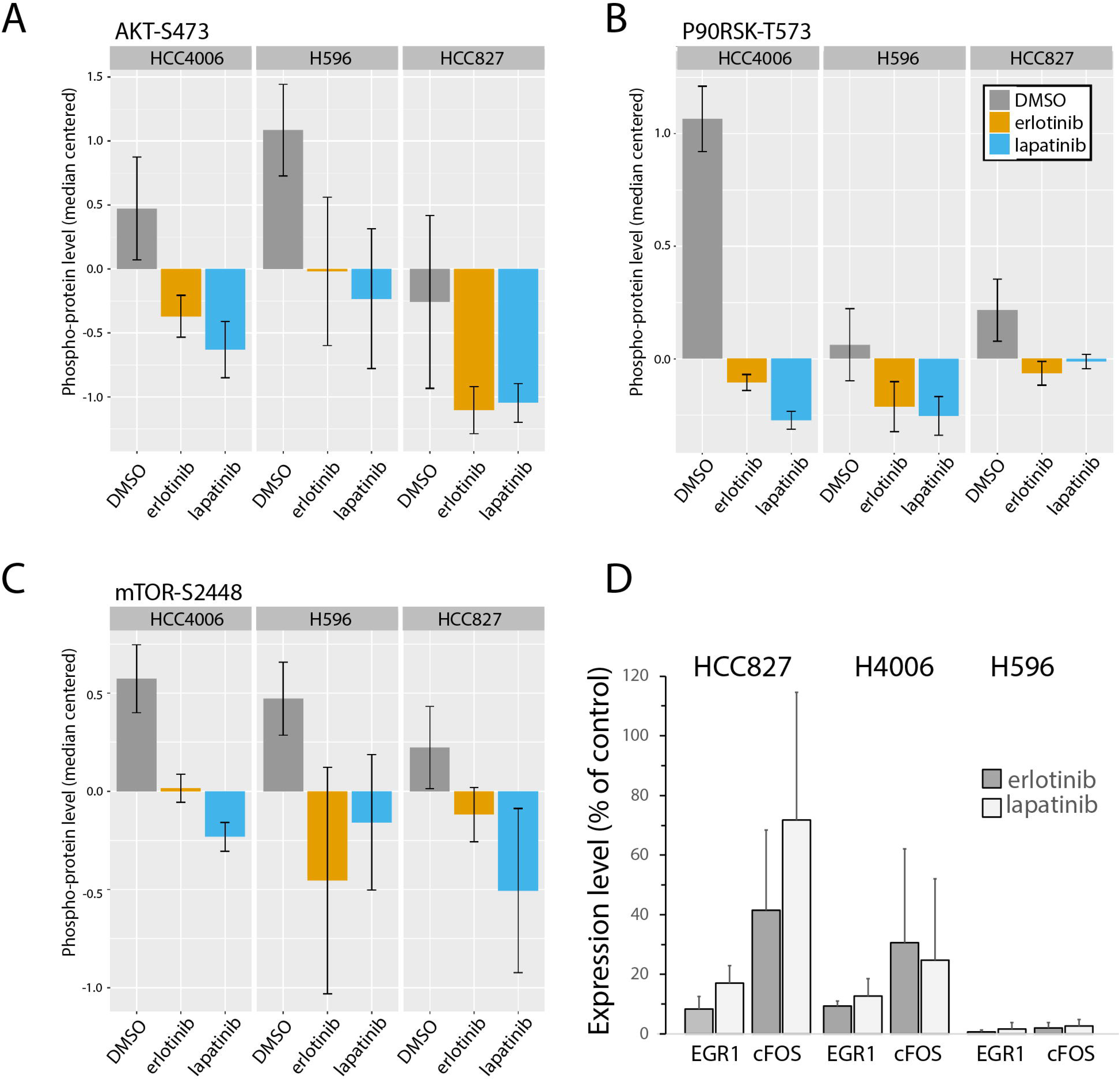
All TKIs effectively inhibit downstream pathway activation. **A-C)** Erlotinib (orange bars) and lapatinib (blue bars) inhibit EGFR pathway activation compared to DMSO control (grey bars). **A)** AKT S473; **B)** mTOR S2448; **C)** P90RSK-T573. Total protein levels of these kinases were not altered (not shown). Data are averages from three independent replicates. **D)** RT-QPCR shows efficacy of inhibitors on EGFR-induced gene expression. Both gefitinib and lapatinib result in a decreased expression of *EGR1* and *cFOS.*Data are shown as the (increase in) the ΔΔCt value relative to the unstimulated control values for these genes. QPCR are averages obtained from 4 independent replicates.

We also performed pull-down assays to examine whether different TKIs differentially affect EGFR protein-protein interactions. Although some inhibitor-specific protein-protein interactions were identified across the various cell lines examined ((HCC827, HCC4006 and HeLa cells expressing EGFR^L858R^, supplementary table 3), no difference that was common between erlotinib/gefitinib with lapatinib was observed. The various TKIs therefore have similar inhibition of EGFR, its pathways and its interactome and therefore do not provide an explanation for the TKI- and mutation-specific intracellular accumulation in EGFR.

### A two-step conformational change model may explain the intracellular accumulation

EGFR is phosphorylated and internalized after its activation by ligand (see e.g. figure 5b and [27]). Once trafficked into early endosomes, it the protein is eventually dephosphorylated and either recycled back to the plasma membrane or transported to the lysosome for degradation. As activated EGFR remaining in the cytoplasm will be recycled back to the membrane, it follows that the inhibition of EGFR activity will result in a (relative) increase in the membrane fraction of the protein. Indeed, quantification of the membrane/cytoplasm ratio of EGFR shows that EGFR-TKIs result in an increased membrane association in cells expressing EGFRwt (figure 7). Interestingly, only lapatinib resulted in this increased membrane association in cells expressing EGFR^L858R^; other TKIs resulted in an increased intracellular accumulation.

**Figure 7:**
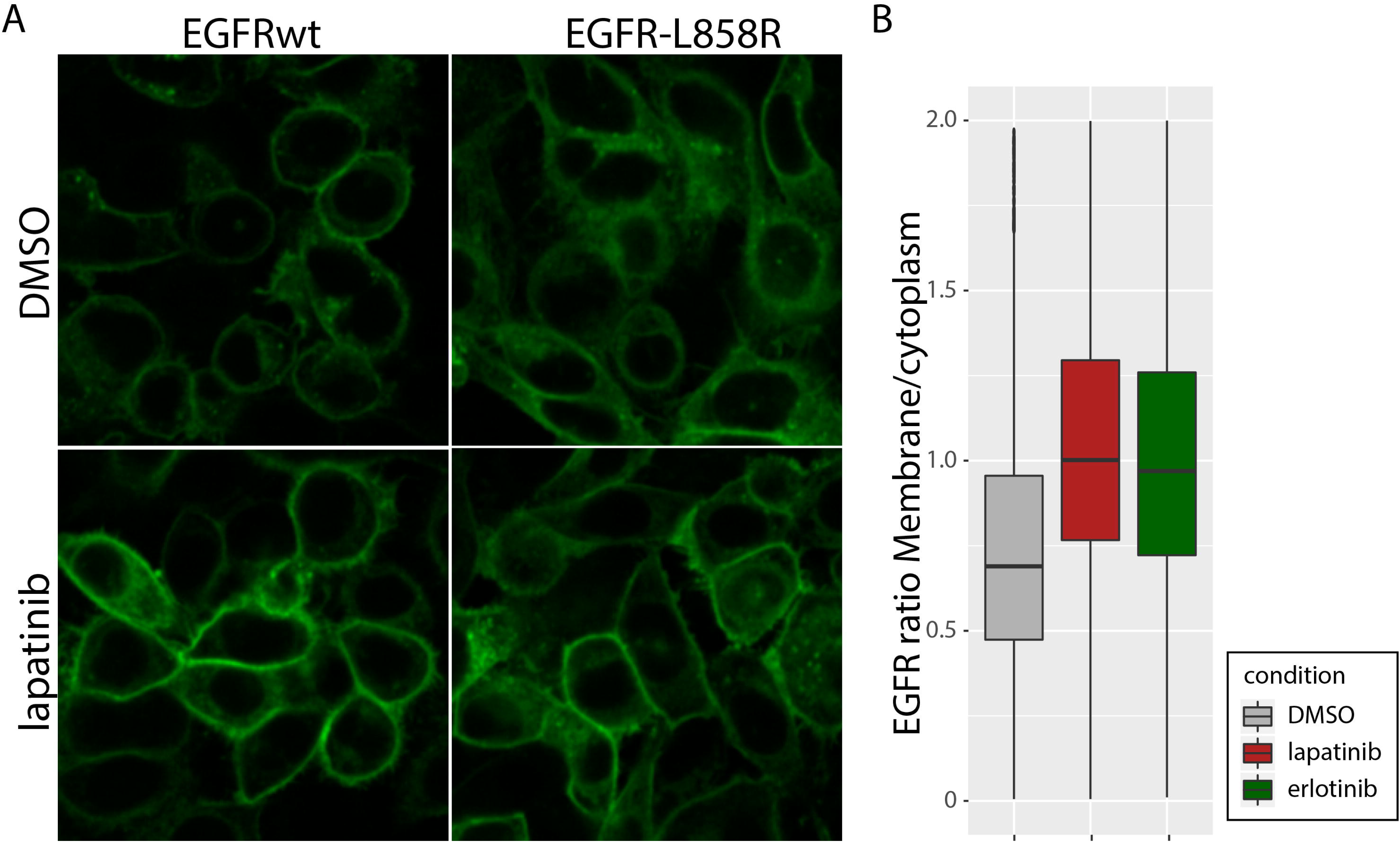
Lapatinib increased membrane association of EGFR. A) example of images showing increased membrane association following treatment with lapatinib in cells expressing EGFRwt or EGFRL858R. B) Quantification of images shown in A.

We hypothesized that the difference between lapatinib and other TKIs on EGFR^L858R^ may lie in the differential conformational preference of TKIs: erlotinib associates with the active conformation while lapatinib traps the protein in an inactive conformation [28-30]. In EGFRwt such conformational preference is TKI-independent: once EGFRwt is dephosphorylated, the protein will adopt an inactive conformation and the protein is recycled to the membrane. However, specific activating mutations such as EGFR^L858R^ destabilize (or even are incompatible with-) the inactive confirmation and promote the protein to adopt its active conformation [28, 29, 31]. Since erlotinib associates with the active conformation it is possible that, in the context of EGFR^L858R^, the TKI remains associated with the protein and this association blocks recycling to the plasma membrane.

To demonstrate clinically effective TKIs remain associated with EGFR^L858R^, we washed out the various inhibitors and monitored intracellular accumulation. The intracellular accumulation indeed depended on the continued presence of the inhibitor (despite EGFR being de-phosphorylated) as removal of competitive inhibitor erlotinib or osimertinib, but not the non-competitive inhibitor dacomitinib, resulted in a reversal the intracellular accumulation in HeLa cells expressing EGFR^L858R^ after >30 minutes of erlotinib/osimertinib withdrawal (figure 8, supplementary figure 6). In lung cancer cell lines harbouring endogenous EGFR mutations, EGFR cannot be re-phosphorylated even after four hours after washout of the inhibitors further confirming that TKIs remain associated with EGFR (supplementary figure 7).

**Figure 8:**
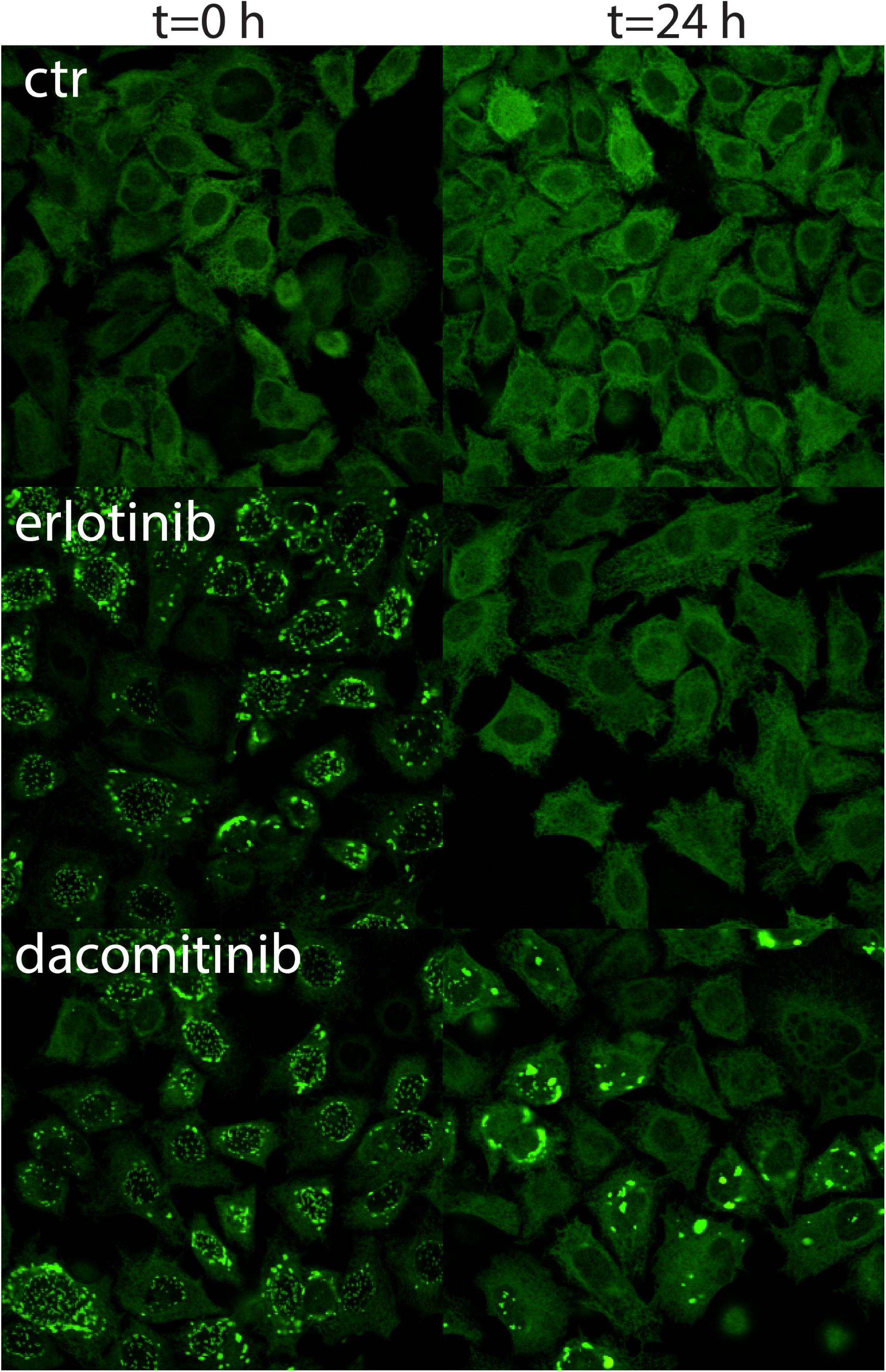
Intracellular accumulation remains dependent on the presence of TKI. Withdrawal of the competitive inhibitor erlotinib (but not the non-competitive inhibitor dacomitinib) reverts the intracellular accumulation demonstrating dependency on TKI presence. Such reversion was also observed following withdrawal or competition of gefitinib and osimertinib (supplementary figure 5).

These results are compatible with the hypothesis that the mutation and TKI-specificity of the intracellular accumulation is be due to two sequential effects: activating mutations firstly lock the protein in an active conformation, TKIs that associate with the active conformation then further affect the conformation of EGFR. Structural studies confirm that TKIs actively affect the conformation of EGFR [28, 29, 31]. This altered conformation then prohibits recycling to the plasma membrane resulting in an intracellular accumulation of the protein.

### Discussion

In this study, we have performed functional analysis on EGFR-mutation constructs to understand why only specific tumor-types respond to EGFR inhibitors, and why only specific inhibitors are clinically effective. We show that the addition of TKIs to cells expressing EGFR-mutation constructs results in a rapid intracellular accumulation of EGFR, but only on mutations that show clinical response to EGFR TKIs and only to EGFR-TKIs that are clinically effective. The accumulation is highly correlated to sensitivity to gefitinib in EGFR-mutated cell lines, and we show that it predicts response to EGFR-TKIs in patients.

Our data has two important clinical implications. First of all, our relatively simple assay can be used to predict the response EGFR TKIs in tumors harboring mutations where this is not yet known. A large database containing the TKI-induced intracellular accumulation all possible EGFR-mutations (alone or in combination with resistance mutations), stably expressed in HeLa cells, would suffice predicting clinical responses, and to which TKI the mutation is likely to be most sensitive. Second, since the intracellular accumulation is seen in cell lines that do not depend on EGFR, our data imply that response to EGFR-TKIs is almost entirely dictated by the type of mutation present, and thus is independent of the cell or tumor type. The tumor type independence of TKI efficacy is supported by several reports where clinical responses to EGFR TKIs have been observed in various (non-pulmonary adenocarcinoma) tumor-types harboring TKI-responsive mutations, [32-36]. This mutation-specificity indicates that all patients with EGFR mutated tumors (regardless of tumor type), that are sensitive to EGFR-TKIs in lung cancer, should be considered for treatment with EGFR-TKIs.

Our data also provides some mechanistic insight into how clinically effective EGFR-TKIs may function: they require two sequential effects on the conformation of the protein. Firstly activating mutations lock the protein in an active conformation. Secondly, TKIs that associate with the active conformation further affect the conformation of EGFR which ultimately prohibits the protein recycling to the plasma membrane. It remains to be determined why the intracellular accumulation results in effective clinical responses. It is possible that intracellular accumulation results in an inactivation of all functions of EGFR, perhaps including those that may not depend on phosphorylation. Such a ‘TKI-induced sequestering of EGFR’ would explain why many (non-pulmonary adenocarcinoma) tumors remain dependent on EGFR for growth, but that inhibition of EGFR-phosphorylation alone is ineffective [37, 38]. If so, targeting EGFR would remain a valid option for tumours that depend on its signalling for growth.

In summary, we provide an assay that can predict whether a tumor harboring an unknown mutation will respond to EGFR-TKIs, and if so, which TKI is most effective. As we show that response to EGFR-TKIs is dictated by the mutation, and not the cell or tumor-type, all patients with sensitive EGFR mutations should, regardless of the type of tumor, be considered for treatment with EGFR-TKIs.

## Methods

### Constructs

EGFR mutation constructs were generated by in-fusion cloning. The backbone of all constructs were essentially as described [24], with eGFP cloned in-frame 3’ to the transmembrane domain. This position was chosen to avoid potential interference with ligand binding or receptor internalization signaling sites. Constructs were cloned into a piggybac vector (System Biosciences, Palo Alto, Ca) allowing for rapid integration using transposase into the host genome. Cell-lines were obtained from the ATCC (Manassas, Virginia). Cells were plated in 96 or 384 well plates for further analysis.

### Image analysis

All images were obtained using an Opera Phenix high-throughput high-content confocal microscope (Perkin Elmer, Hamburg, Germany). At least 10 images were obtained per well so that an experiment involving a single construct, 6 conditions (5 inhibitors + control) at 10 different dilutions typically would produce >600 images per timepoint. Channels were independently excited to minimize potential spectral overlap. Image analysis was performed in bulk using Harmony software (Perkin Elmer) using identical settings within each experiment. Èxperiments described in current manuscript were performed at least in two independent replicates. Data was further analysed using R.

### Stainings

EGFR antibody (clone H11, DAKO, Amstelveen, the Netherlands) and a phospho-specific EGFR antibody (AB32430, anti phospho Y1068, Abcam, Cambridge, UK) were used at 1:500 dilution for both western blot and immunohistochemistry. Secondary antibodies used were alexafluor 647 goat anti-mouse (A21240, Invitrogen, Bleiswijk, the Netherlands) and alexafluor 488 goat anti-rabbit (A11008, Invitrogen, Bleiswijk, the Netherlands). Hoechst and WGA were used as counterstain to visualize nucleus and membranes respectively.

### RT-QPCR

RNA was extracted from cells using the RNeasy mini kit (Qiagen, Venlo, the Netherlands). RT-QPCR was performed using Taqman probes (Applied Biosystems, Bleiswijk the Netherlands) according to the manufacturers’ instructions. Expression levels of *cFOS* and *EGR1* were evaluated relative to *POP4* and *GAPDH* controls.

### Patients

We identified pulmonary adenocarcinoma patients harbouring EGFR mutations from routine diagnostics within the Erasmus MC. For patients screened in 2016, no selection was made other than presence of a mutation in the *EGFR* gene. The data was further expanded with patients screened in 2017 and 2018 but not including patients with exon 19 deletions or the L858R missense mutation (thus selecting for rare mutations). We generated constructs for these mutations. If multiple mutations were identified, the prediction of response was made based on the one with highest IC50. Response predictions were performed with the experimenter blinded to the clinical outcome. The separation into responders/non-responders was performed blinded to clinical outcome using a predefined cutoff of 500 nM. This cutoff was chosen prior to the analysis and was based on maximal concentrations of inhibitor that are achieved in patients, though there is a large inter patient variability [16]. Progression free survival was defined as the time to progression to first line TKI treatment. Patients were censored in case of enduring clinical response or when lost to follow-up.

### RPPA

All samples were prepared according to the guidelines of the MD Anderson functional proteomics RPPA core facility, where all RPPA experiments were subsequently run. Cells were maintained under normal (serum supplemented) culture conditions and inhibitors or DMSO were added two hours prior to cell lysis. RPPA experiments were generated in three experiments, with each experiment performed in a separate week at a different cell-passage number to ensure complete independence.

## Supplementary Materials

supplementary figure 1: Erlotinib rapidly induces intracellular accumulation of EGFR.

supplementary figure 2: Intracellular EGFR accumulation in U87 cells.

supplementary figure 3: Intracellular accumulation of various mutation constructs.

supplementary figure 4: All TKIs effectively inhibit EGFR phosphorylation.

supplementary figure 5: Intracellular accumulation consists of dephosphorylated EGFR.

supplementary figure 6: Intracellular accumulation remains dependent on the presence of TKI.

supplementary figure 7: Rephosphorylation after washout of EGFR TKIs

Supplementary movie 1: Erlotinib rapidly induces intracellular accumulation of EGFR.

## Acknowledgments

This work was supported by a grant from KWF kankerbestrijding, grant number 11125.

## Author contributions

Conceptualization, PJF; Methodology, P.J.F, M.v.R., J.A. and P.S.S.; Investigation, Y.G., M.d.W., I.d.H. and B.V.; Writing – Original Draft, P.J.F; Writing – Review & Editing Y.G., M.d.W., D.M, I.d.H., B.V., M.v.R, J.A and P.S.S.; Funding Acquisition, P.J.F. and P.S.S.; Resources, M.v.R. and D.M.; Supervision, P.J.F, J.A. and P.S.S.

**Supplementary movie 1: Erlotinib rapidly induces intracellular accumulation of EGFR.** Within three minutes of the addition of erlotinib, intracellular accumulation is visible in the peri-nuclear region of the cell. Images are taken every 20 seconds, total timeframe of the movie is six minutes.

**supplementary figure 1: Erlotinib rapidly induces intracellular accumulation of EGFR.** Within 3 minutes after the addition of inhibitor, intracellular EGFR accumulation is observed. Images are stills from supplementary movie 1, taken at 1.5 minute intervals.

**supplementary figure 2:: Intracellular EGFR accumulation in U87 cells.** A) Similar to observed in HeLa cells, U87 also respond to clinically effective EGFR TKIs by the intracellular accumulation of EGFR. B) quantification of images in A). Dacomitinib is toxic to U87 cells and therefore is not included in this analysis. In U87 cells EGFR accumulates at a slower pace than in HeLa cells (24+ hrs) with only little accumulation visible at 2 hr. Despite these temporal and quantitative differences, our results demonstrate that the intracellular accumulation occurs in other cell-lines, but only those harboring activating EGFR mutations.

**supplementary figure 3: Intracellular accumulation of various mutation constructs.** Constructs EGFR^L858R^, EGFR^E746_A750del^, EGFR^L747_E749del^, EGFR^S768I^, and EGFR^G719S^ all responded to 10 μM erlotinib or gefitinib by rapidly inducing EGFR accimulation. None of the other mutation constructs showed such accumulation.

**supplementary figure 4: All TKIs effectively inhibit EGFR phosphorylation.** A) Western blots showing EGFR phosphorylation under normal culture conditions in the H1975, H596 and H460 cell lines. H1975 cells harbor the T790M resistance mutation and only 3^rd^ generation TKI osimertinib inhibits EGFR phosphorylation. No phosphorylation was observed in H596 and H460 cells. For reference, a lane of HCC827 cells is included. B) Quantification of the phospho-EGFR signal using image analysis. In unstimulated cells (top panels), EGFR is phosphorylated in HCC827 cells, and the addition of any TKI results in a rapid dephosphorylation. Little phospho-EGFR was detected in NCI-H1975, and no virtually no phospho-EGFR was detected in NCI-H460 or NCI-H596 cells (in line with our western blot data). EGF stimulation (bottom panels) results in a pronounced increase in phospho-EGFR staining in NCI-H1975, NCI-H460 or NCI-H596 cells and the addition of TKI results in a rapid dephosphorylation. Virtually identical data was obtained in three independent replicate experiments (not shown).

**supplementary figure 5: Intracellular accumulation consists of dephosphorylated EGFR.** The accumulation in erlotinib treated HeLa cells expressing EGFR^L858R^, do not stain positive when using a phospho-specific antibody (left panel, staining in red) but stain positive when using a pan-EGFR antibody (right panel).

**supplementary figure 6: Intracellular accumulation remains dependent on the presence of TKI.** Intracellular accumulation remains dependent on the presence of TKI, since withdrawal of the inhibitors erlotinib or osimertinib (but not the non-competitive inhibitor dacomitinib) reverts the accumulation. The accumulation remained stable after withdrawal of the competitive inhibitor gefitinib, but these could be competed out by co-application of GW2974, a competitive EGFR inhibitor and analogue of lapatinib. This inhibitor was identified as part of a screen using the Lopac1280 library to identify compounds that prevent gefitinib induced intracellular accumulation). Of note, GW2974 is a structural analogue of lapatinib [39].

**supplementary figure 7: Effect of TKIs after washout.** In the HCC827 lung cancer cell line harbouring an endogenous EGFR mutations, EGFR cannot be re-phosphorylated even after four hours after washout of the inhibitors. A) example of phosphor-EGFR staining following EGF stimulation in the presence of EGFR TKIs. B) Quantification of images in A at various timepoints after washout (0, 15, 60 and 240 minutes).

