## Supplementary figures and images for "Effective Tyrosine Kinase Inhibitors result in the intracellular accumulation of EGFR and allows response prediction in patients"

### Supplementary figure 1

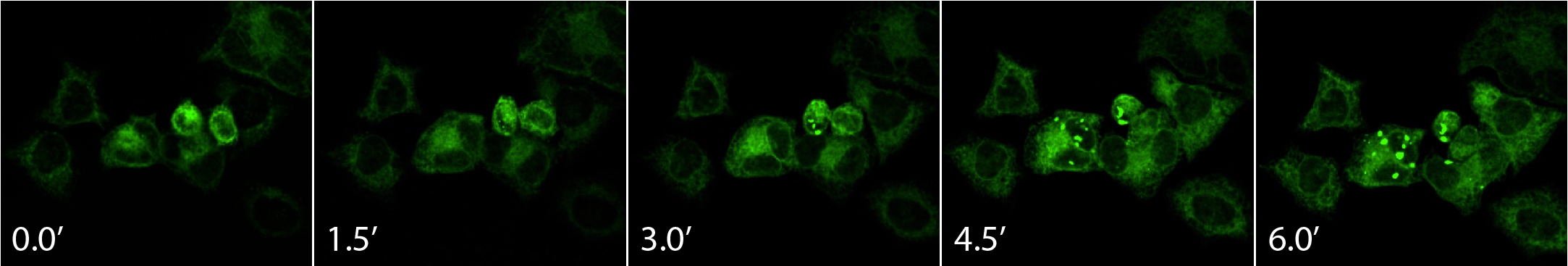

### Supplementary figure 2

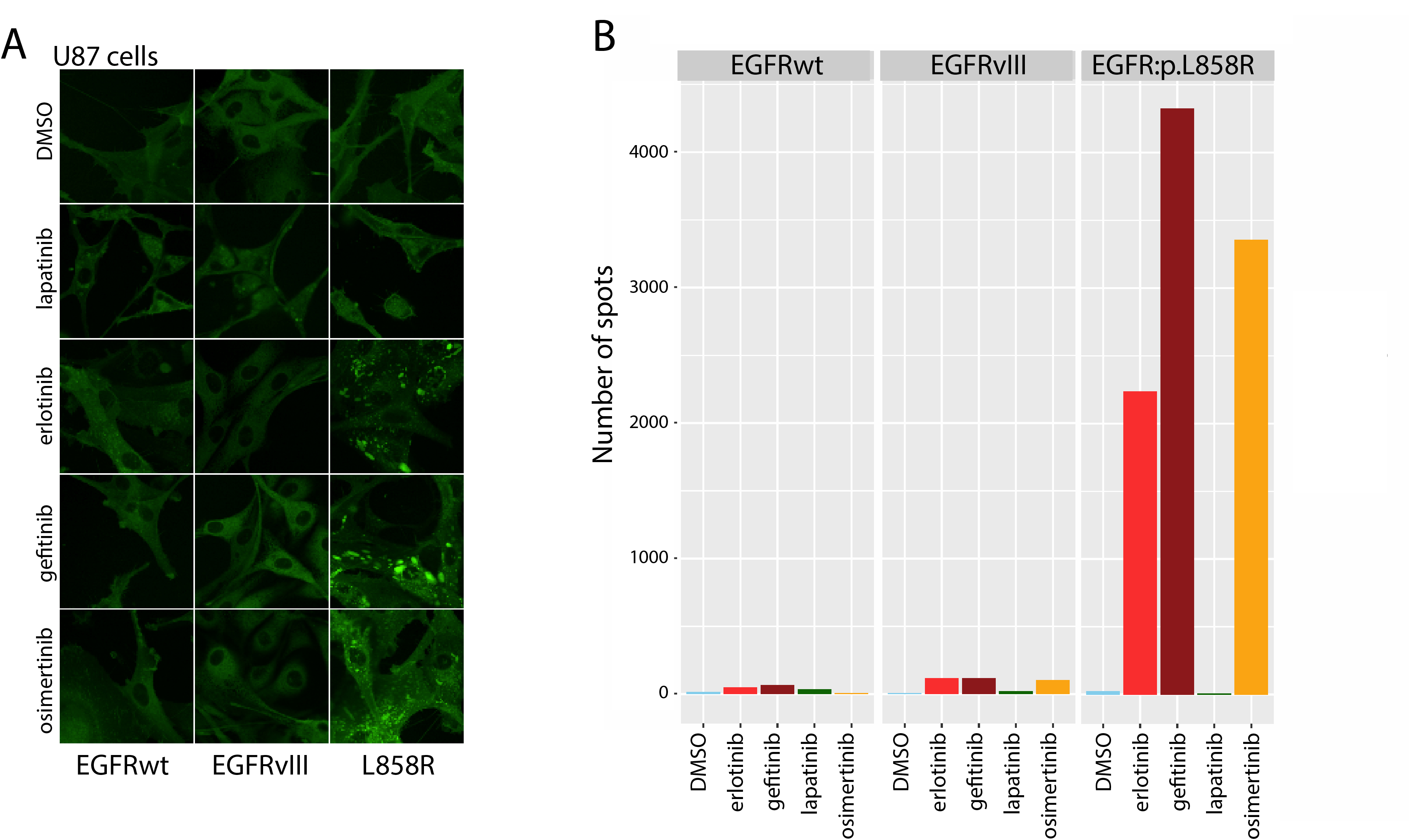

### Supplementary figure 3

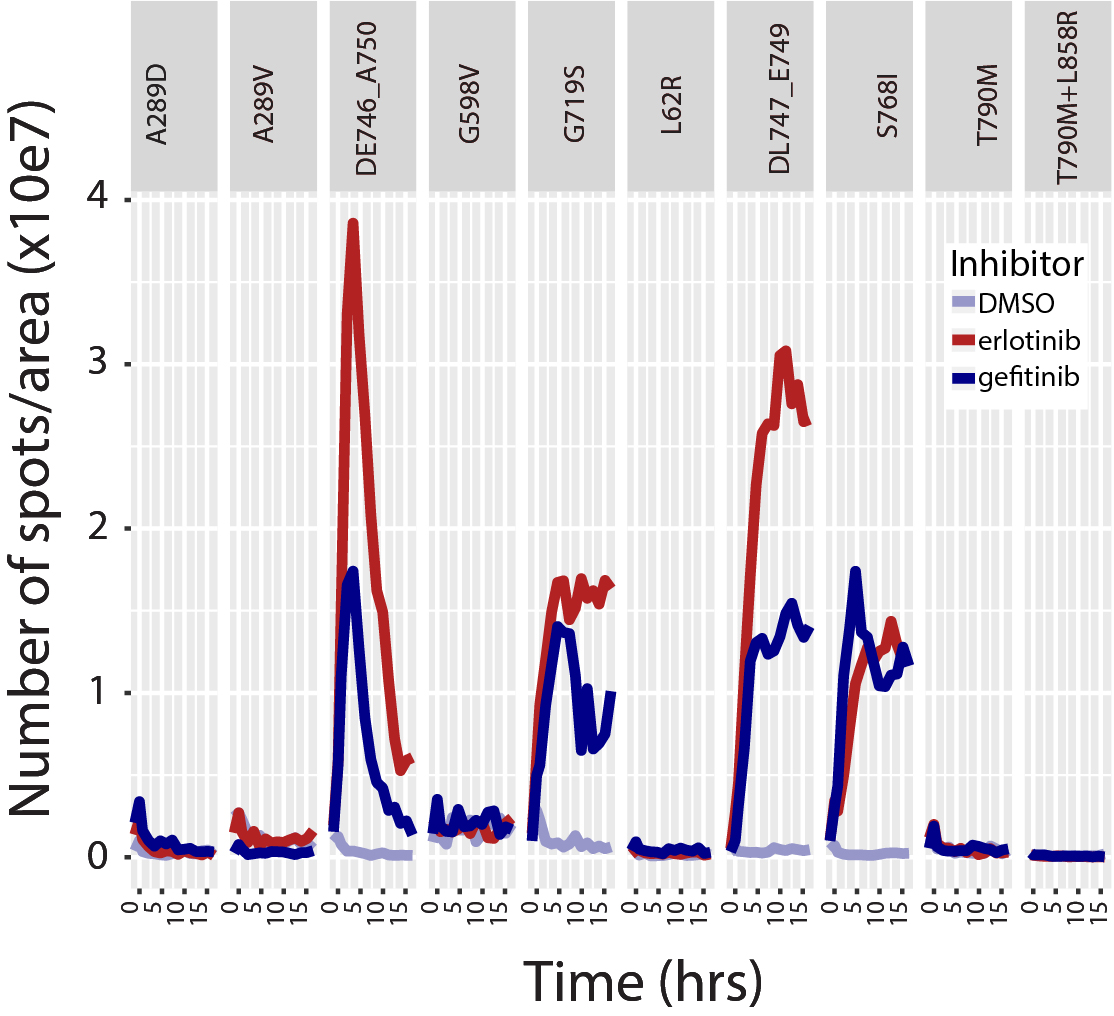

### Supplementary figure 4

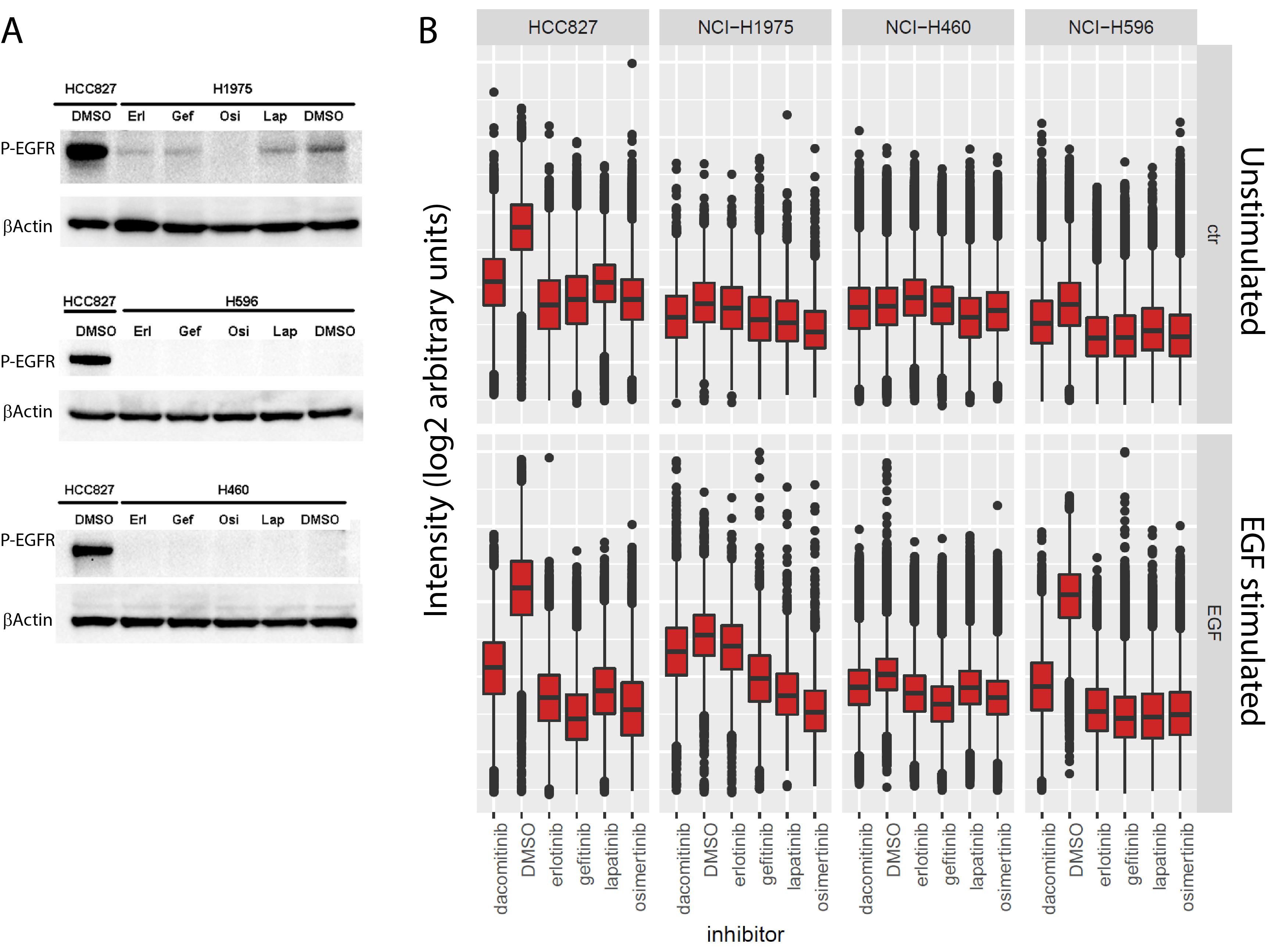

### Supplementary figure 5

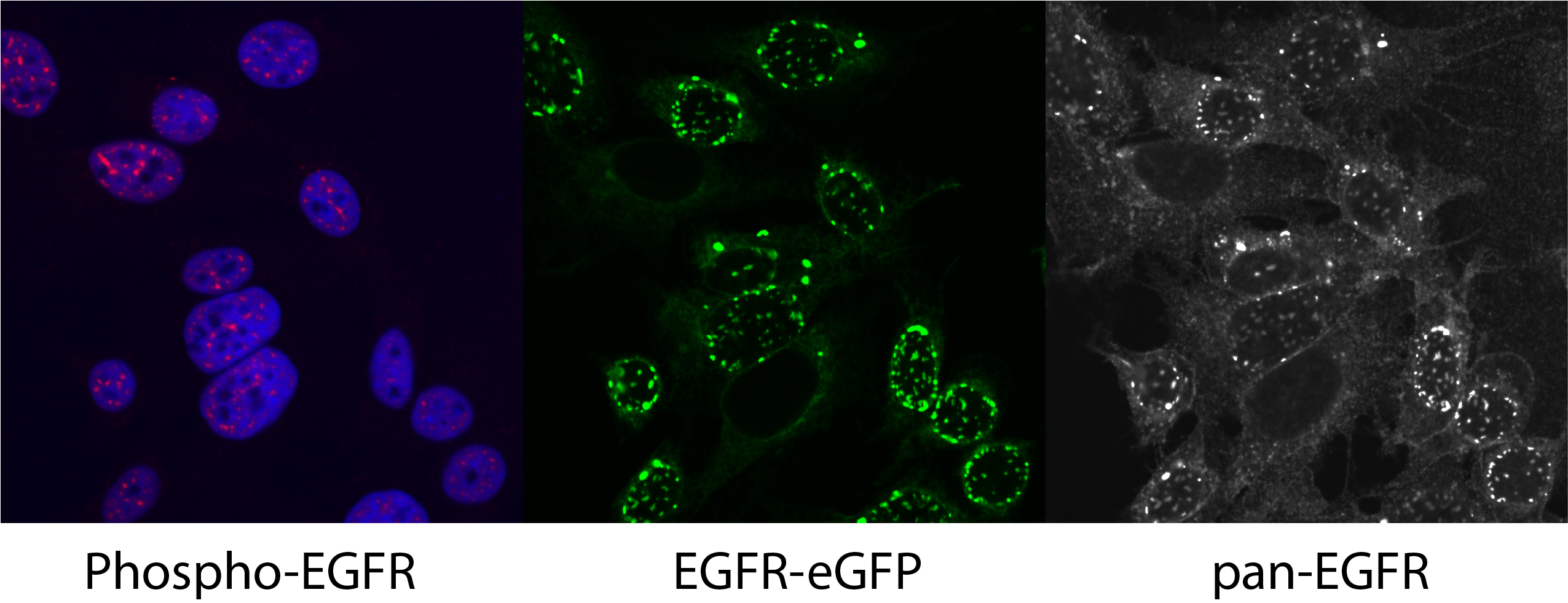

### Supplementary figure 6

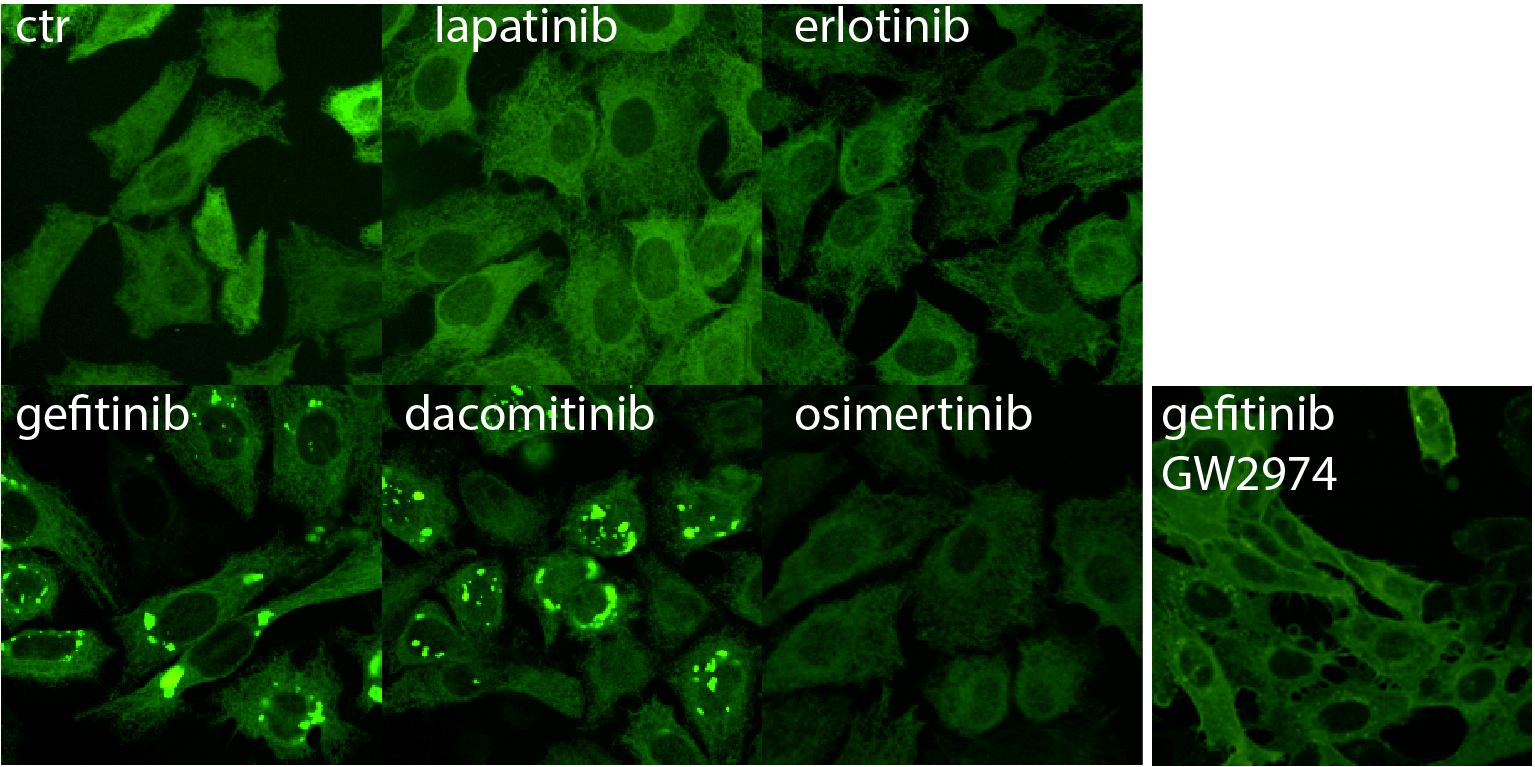

### Supplementary figure 7

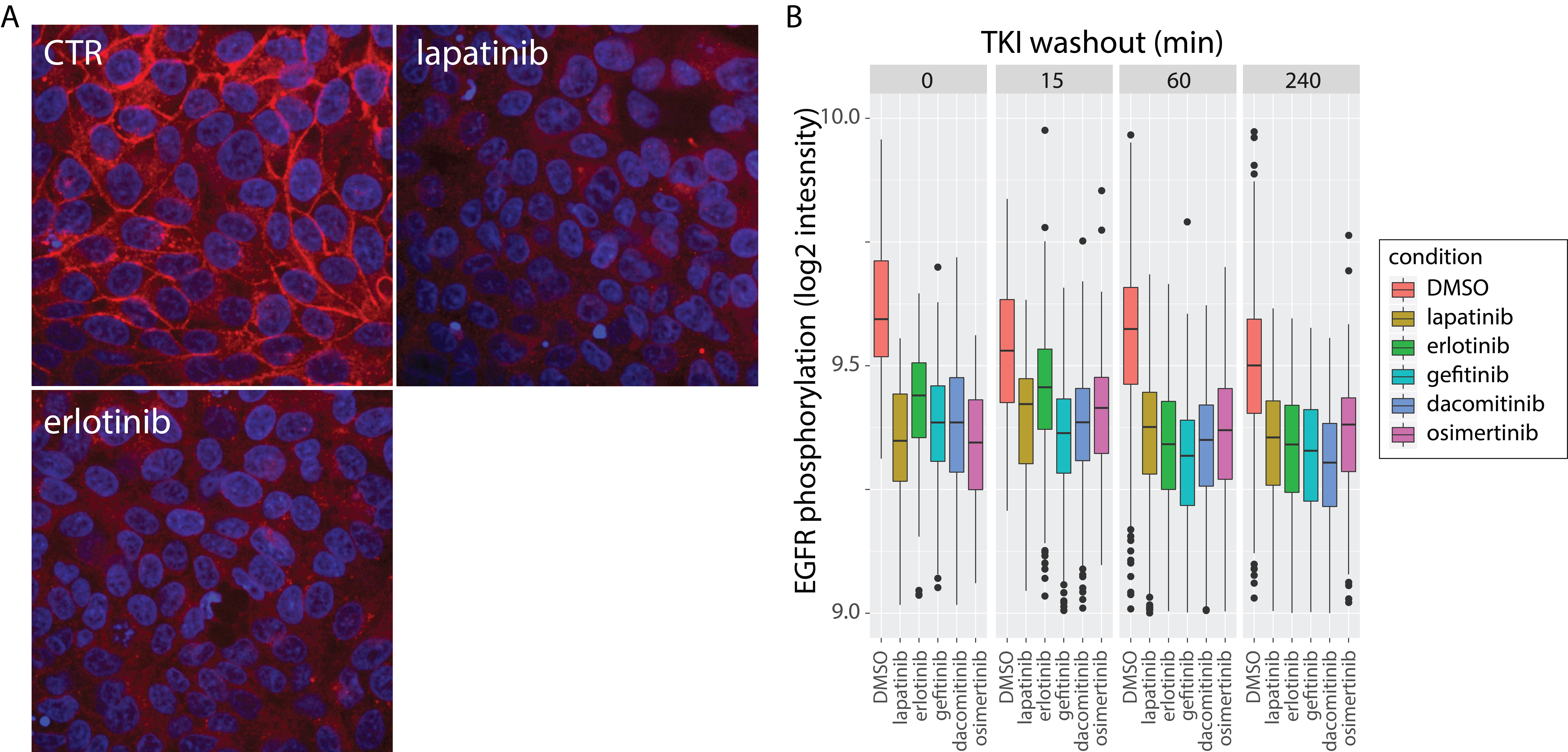
